# *AGAMOUS-like 6* and *MYB DOMAIN PROTEIN 80* regulate the development of female and male cones in *Pinus densiflora* S. et Z

**DOI:** 10.1101/2023.10.18.562949

**Authors:** Dayoung Lee, Yang-Gil Kim, Kyu-Suk Kang

## Abstract

To shorten the long breeding period of forest trees, it is necessary to explore biological factors and key genes involved in reproduction. We compared the transcriptomes of female and male cones and then identified major biological processes and hub genes associated with the development of female and male cones in *Pinus densiflora*. The reproductive and vegetative tissues of *P. densiflora* were sampled from various developmental stages and a reference transcriptome was generated. Functional annotation and Gene Ontology enrichment analysis of differentially expressed genes were performed as well as gene co-expression network analysis for identifying the hub genes involved in the cone development. Lignin biosynthesis and cell wall loosening were activated in the early and late female cones, respectively. Metabolism of pollen wall components was predominant in the early male cones compared with the later stage. In addition, phytohormone-related genes were highly expressed in the early female cones in contrast to the early male cones. Finally, we found that *AGAMOUS-like 6* and *MYB DOMAIN PROTEIN 80* were hub genes of the female and male cone development, respectively. This study suggests that various biological factors contribute to reproductive developmental processes. Furthermore, these data will provide the new insight into biotechnology and breeding methods for improving cone and seed production of *P. densiflora*.

## Introduction

For seed plants, the process of transition from vegetative to reproductive phase is essential for survival. In the case of angiosperms such as *Arabidopsis*, many genetic studies on reproductive development have been conducted, but studies on the function or expression of flowering genes in perennial conifers, which are gymnosperms, are relatively insufficient (Ma et al., 2021). Conifers have a long reproductive period, large and complex genomes. The lack of reliable reference genomes makes it difficult to understand the gene function of conifers. Moreover, because mutations rarely occur in gymnosperm species, the functions of many genes remain unknown (Moyroud et al., 2010). The genome sizes of *Pinus* genus are quite large, and there are few species whose whole genome has been analyzed in *Pinus*. Therefore, studies for genetically understanding complex phenomena such as flowering have been limited in *Pinus*.

Perennial trees have a long vegetative growth period before reproductive growth begins, and unlike angiosperms, which take one to two years to flower, conifers generally take three years to produce complete cones (Singh, 1978). Due to these characteristics, conifer trees generally have a long breeding period, so it takes decades to develop new varieties by traditional breeding methods such as selective breeding and hybrid breeding. As such, it is difficult for conifer with slow growth and long breeding period to adequately respond to the rapidly changing demands of modern society. Therefore, if improved varieties are developed by, for example, regulating the expression of genes involves in flowering and inducing early cone development, progeny tests will be advanced, and the overall yield of cones will be increased.

The processes involved in flower formation are conserved in angiosperm and gymnosperm species of seed plants. First, the beginning of flowers requires the passage from the vegetative phase, when shoots and leaves are generated, to the reproductive phase (Mouradov et al., 1999). Then, flower meristem identity genes promote flower development. *MADS-box* transcription factor genes are floral organ identity genes and largely conserved between gymnosperms and angiosperms (Scutt et al., 2006). However, the flowering in plants does not appear only through the expression of specific flowering genes, but through numerous biological processes and environmental influences, such as various phytohormones and metabolic actions. Therefore, to understand the flowering phenomenon of conifer trees, it is necessary to explore the entire biological processes that occur during the cone developmental period.

*Pinus densiflora* Sieb. & Zucc. is well adapted to dry and barren environments and can survive in a wide range. It occupies the largest area as a single tree species in South Korea and inhabits parts of China and Japan. *P. densiflora* has been used as wood for a long time, and genetic studies related to this species have been conducted mainly on wood and cone productivity. Pine trees have a long juvenile phase and generally take more than 5 to 10 years to produce cones. Therefore, it is important to shorten the juvenile phase to enable efficient and fast tree breeding.

Several studies have been conducted to explore environmental factors that cause mass flowering using gene expression analyses. Nitrogen was identified as a key regulator of flowering in *Fagus crenata* (Miyazaki et al., 2014) based on the expression analysis of flowering genes. In addition, drought was reported as a trigger for flowering of *Shorea beccariana* in the study using genome-wide expression data (Kobayashi et al., 2013). Marker-assisted selection was also performed to apply it to fast breeding, which shortens the juvenile phase of apple (*Malus* × *domestica*) (Flachowsky et al., 2011). And in Chickpea (*Cicer arietinum*), which belongs to Fabaceae, there is a study to develop genetic marker by exploring candidates involved in early flowering and seed size (Manchikatla et al., 2021).

In this study, we compared differentially expressed gene expression in various tissues and then identified major biological processes and hub genes associated with the reproductive development of *P. densiflora* to obtain biological factors that play a key role in female and male cones. Understanding the biological processes of reproductive development in conifer could help us to understand the molecular regulation of cone development by investigating differential expression of flower-related genes in cones (Mathews and Kramer, 2012). This study shows the possibility of the molecular regulation of cone development and of controlling initiation time of the reproductive phase in *Pinus* conifer species.

## Results

### Differential expression of reproductive and vegetative tissues

Principal Component Analysis (PCA) and hierarchical cluster analysis using Pearson correlation were conducted to identify the differential expression among library samples. Both analyses identified tissue-specific clusters, indicating the statistical significance of biological replicates (Supplemental Fig. S1). Hierarchical cluster analysis using log_2_ normalized gene expression values indicated that the total samples were clustered into several groups: apex and female cones clustered into a very similar group, whereas needles and stems were classified into the other group.

Female cones were not completely clustered at each stage of the developmental phase, and this is because the rate of female cone development differed among the biological replicates. Male cone samples on the late developmental stage were separated from other stages of male cones because the last stage was sampled in a state where all the pollen was blown. The correlation analysis between the female and male cones yielded two clearly separated clusters, confirming distinct expressions of sex-specific genes.

Using threshold values of FC ≥ 4 in the gene expression difference and *p*-value < 0.001, 25307 genes were identified as DEGs in the total 45 pairwise comparisons between any two of the tissue samples (Supplemental Fig. S2). The comparison of root and other tissues showed the most DEGs, and the comparison of apex and female cones showed the least difference in gene expression.

Especially, we further identified a subset of 6368 DEGs expressed in five different tissues (A, F1, F3, M1, M2): 1623, 597, 4212 and 1343 DEGs in A vs F1, F1 vs F3, F1 vs M1 and M1 vs M2 comparisons, respectively (Fig. 1). We found 4212 genes showed sex-specific expression in early female and male cones, and almost the same number of genes were up-regulated both in the female and male cones (2104 in female cones and 2108 in male cones, respectively). These five different tissues separated into two distinct clusters based on the expression difference (Fig. 2): female cones at developmental stages F1 and F3 grouped together with apex (A); male cones at stages M1 and M2 grouped together as the separate cluster. This clustering suggested that the transcriptome of early apex is more like female cones than male cones, and that early female cones are more similar with apex than late female cones.

**Figure 1.**
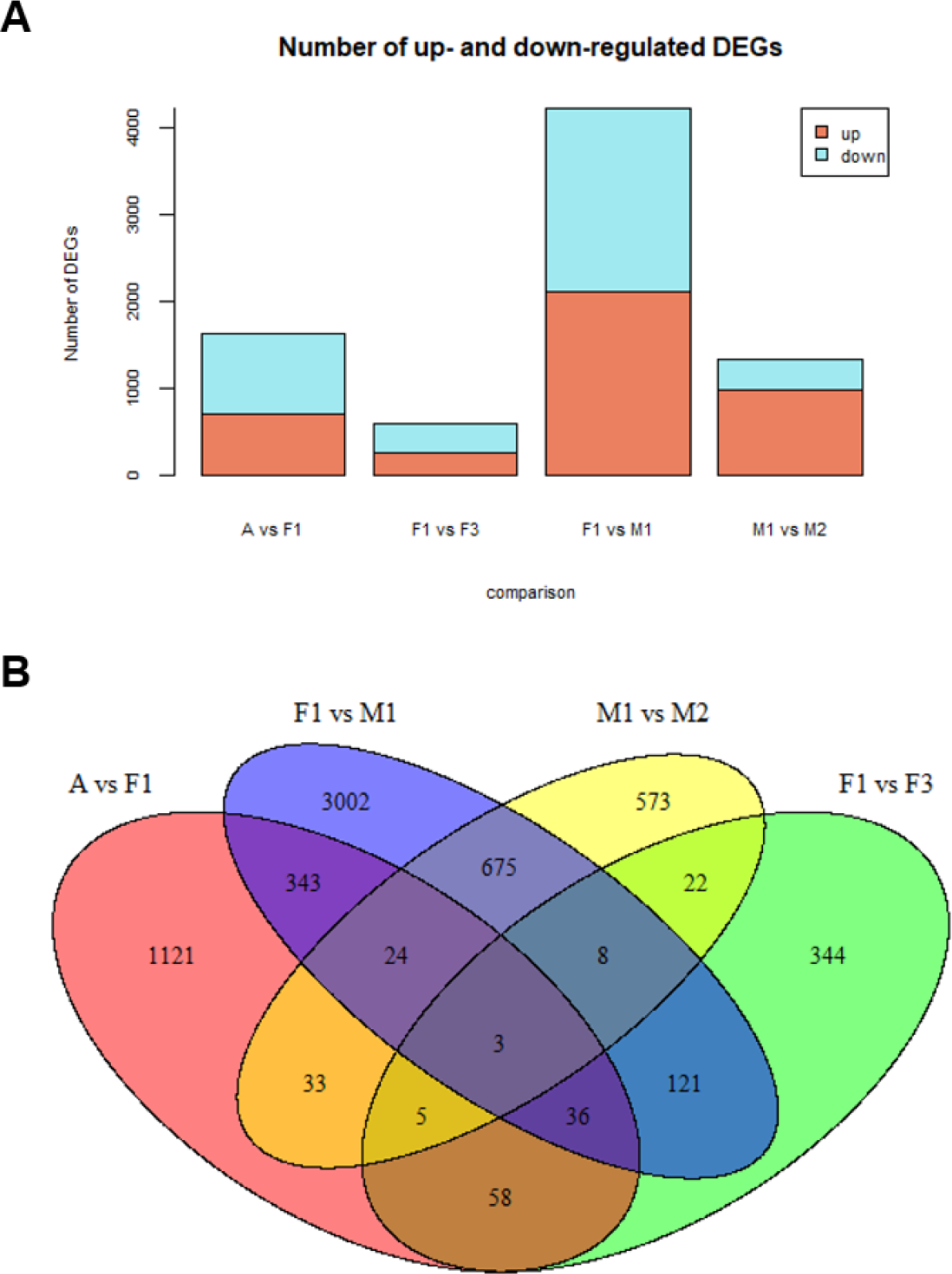
The number of Differentially Expressed Genes (DEGs) in each comparison. **A)** Histogram of up/down-regulated DEGs. **B)** Venn diagram indicating the number of sharing DEGs.

**Figure 2.**
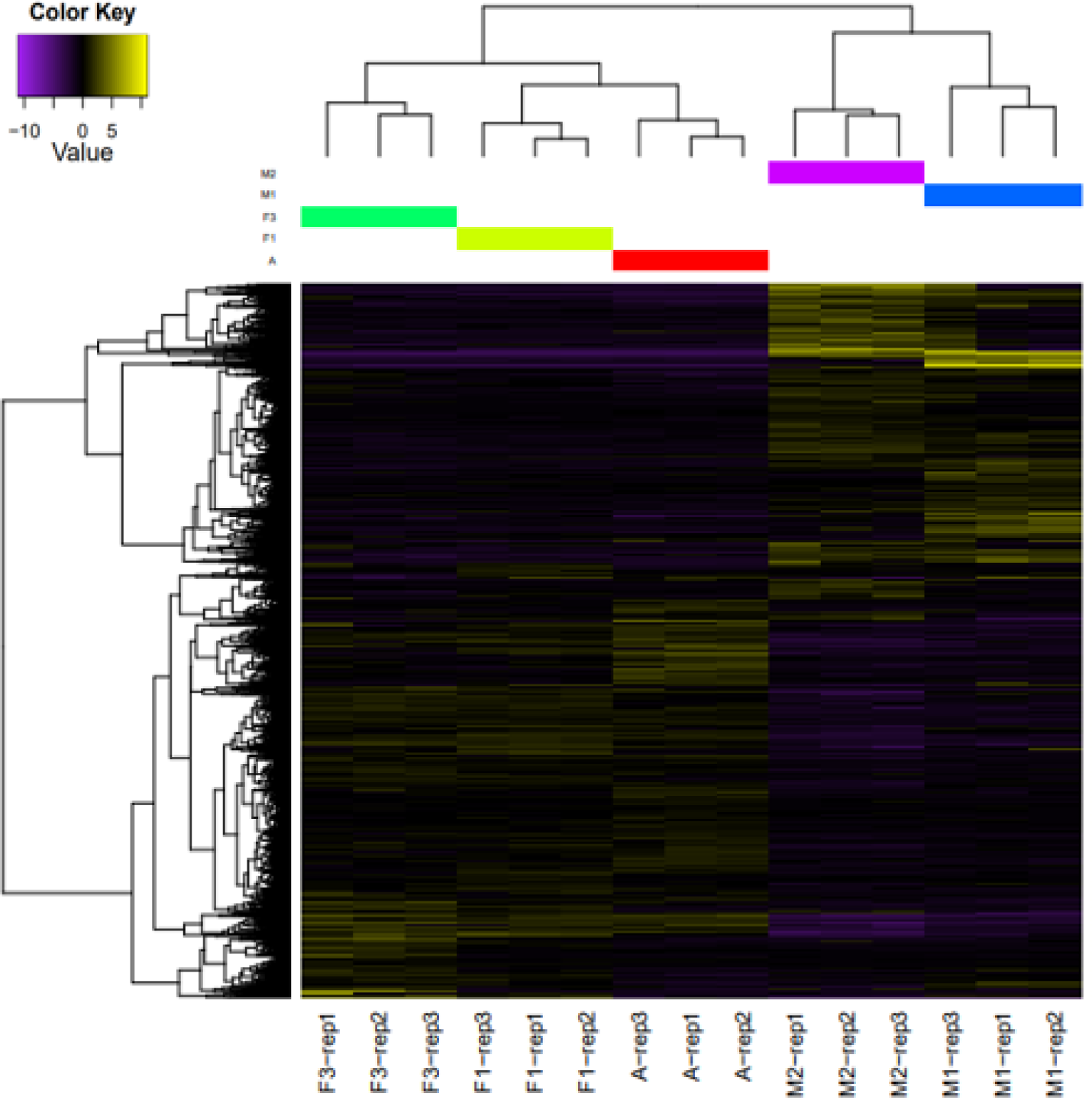
Heatmap of differentially expressed genes of apex (A), female cones (F1 and F3), and male cones (M1 and M2). Female cones include early and late developmental stages and male cones include early and mid developmental stages.

### Biological processes differ according to the cone developmental stage

Functional enrichment analyses were conducted using the DEGs found in the female and male cones at two different developmental stages (F1 and F3; M1 and M2, respectively). GO enrichment analyses identified different patterns of biological processes among the four different tissues of *P. densiflora* (Table 1). In F1 vs F3, ‘lignin biosynthetic process (GO:0009809)’ was significantly enriched in early female cones and ‘plant-type cell wall loosening (GO:0009828)’ was enriched in late female cones. In M1 vs M2, ‘lipid catabolic process (GO:0016042)’, ‘syncytium formation (GO:0006949)’, ‘lignin biosynthetic process (GO:0009809)’ and ‘pectin catabolic process (GO:0045490)’ were significantly enriched in early male cones. There was no significant GO term in mid male cones when compared with early male cones.

**Table 1.**
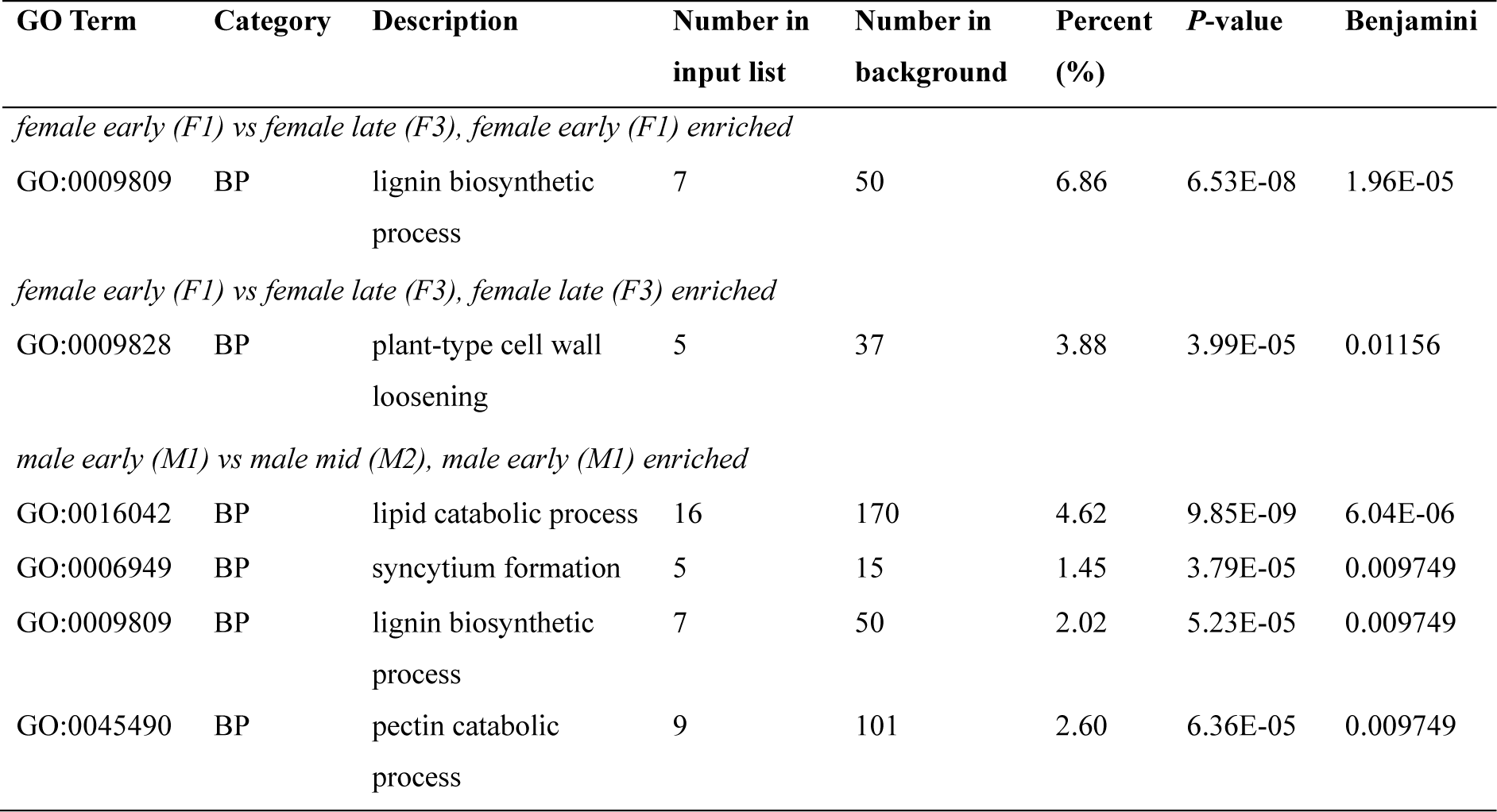
Gene Ontology enrichment results on the different developmental stages of the female and male cones of *Pinus densiflora*.

| GO Term | Category | Description | Number in input list | Number in background | Percent (%) | P-value | Benjamini |
| --- | --- | --- | --- | --- | --- | --- | --- |
| *female early (F1) vs female late (F3), female early (F1) enriched* |  |  |  |  |  |  |  |
| GO:0009809 | BP | lignin biosynthetic process | 7 | 50 | 6.86 | 6.53E-08 | 1.96E-05 |
| *female early (F1) vs female late (F3), female late (F3) enriched* |  |  |  |  |  |  |  |
| GO:0009828 | BP | plant-type cell wall loosening | 5 | 37 | 3.88 | 3.99E-05 | 0.01156 |
| *male early (M1) vs male mid (M2), male early (M1) enriched* |  |  |  |  |  |  |  |
| GO:0016042 | BP | lipid catabolic process | 16 | 170 | 4.62 | 9.85E-09 | 6.04E-06 |
| GO:0006949 | BP | syncytium formation | 5 | 15 | 1.45 | 3.79E-05 | 0.009749 |
| GO:0009809 | BP | lignin biosynthetic process | 7 | 50 | 2.02 | 5.23E-05 | 0.009749 |
| GO:0045490 | BP | pectin catabolic process | 9 | 101 | 2.60 | 6.36E-05 | 0.009749 |

### Identification of sex-specific biological processes between the early female and male cones

Functional enrichment analyses were conducted with the DEGs between the early female and male cones and we identified specific biological processes in each of the early cone tissues of *P. densiflora* (Fig. 3). In the early female cones (F1), GO terms related to plant hormones such as ‘response to gibberellin (GO:0009739)’, ‘gibberellic acid mediated signaling pathway (GO:0009740)’ and ‘response to auxin (GO:0009733)’ were up-regulated compared to M1. Flowering related terms such as ‘anther development (GO:0048653)’ and ‘flower development (GO:0009908)’, and cell wall related terms such as ‘pectin catabolic process (GO:0045490)’, ‘lignin biosynthetic process (GO:0009809)’ and ‘plant-type cell wall modification involved in multidimensional cell growth (GO:0009831)’ were significantly enriched in F1.

**Figure 3.**
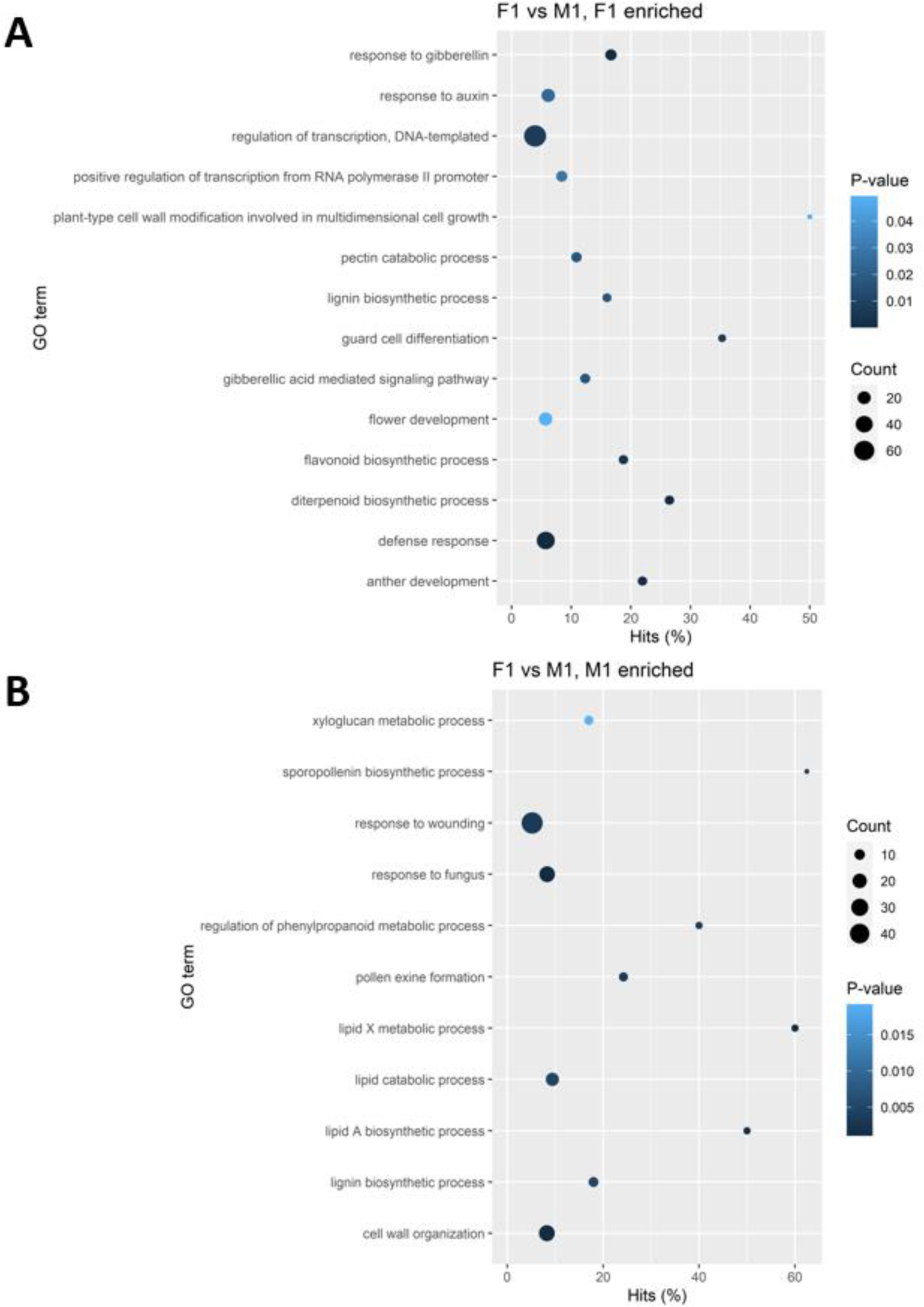
Gene Ontology (GO) enrichment analysis on the differentially expressed genes between the early female and male cones of *Pinus densiflora*. **A)** Female cone enriched and **B)** male cone enriched biological GO terms.

On the other hand, in the early male cones (M1), GO terms related to lipid metabolism such as ‘lipid X metabolic process (GO:2001289)’, ‘lipid A biosynthetic process (GO:0009245)’ and ‘lipid catabolic process (GO:0016042)’ were up-regulated compared to F1. Cell wall related terms such as ‘cell wall organization (GO:0071555)’, ‘lignin biosynthetic process (GO:0009809)’ and ‘xyloglucan metabolic process (GO:0010411)’ were also significantly enriched in M1.

### Identification of hub genes specific to the development of female and male cones

A total of 19 modules were identified and then labeled with different colors and shown as a hierarchical clustering dendrogram (Supplemental Fig. S3). The relationship between modules and traits is shown in Supplemental Fig. S4. In the Fig. 4, the line graph matches the correlation heatmap in Supplemental Fig. S4 where ‘tan’ module eigengenes are higher in female cone groups and ‘yellow’ module eigengenes are higher in the male cone groups. According to expression levels of clustered genes in each module, tan and yellow modules were selected for co-expression analysis and identification of hub genes. GO enrichment analysis was performed on the eigengenes in tan and yellow modules (Supplemental Table S1). The tan module was enriched with genes involved in the floral meristem determinacy and defense response. The yellow module was related to the pollen formation, cell wall organization and lipid metabolism.

**Figure 4.**
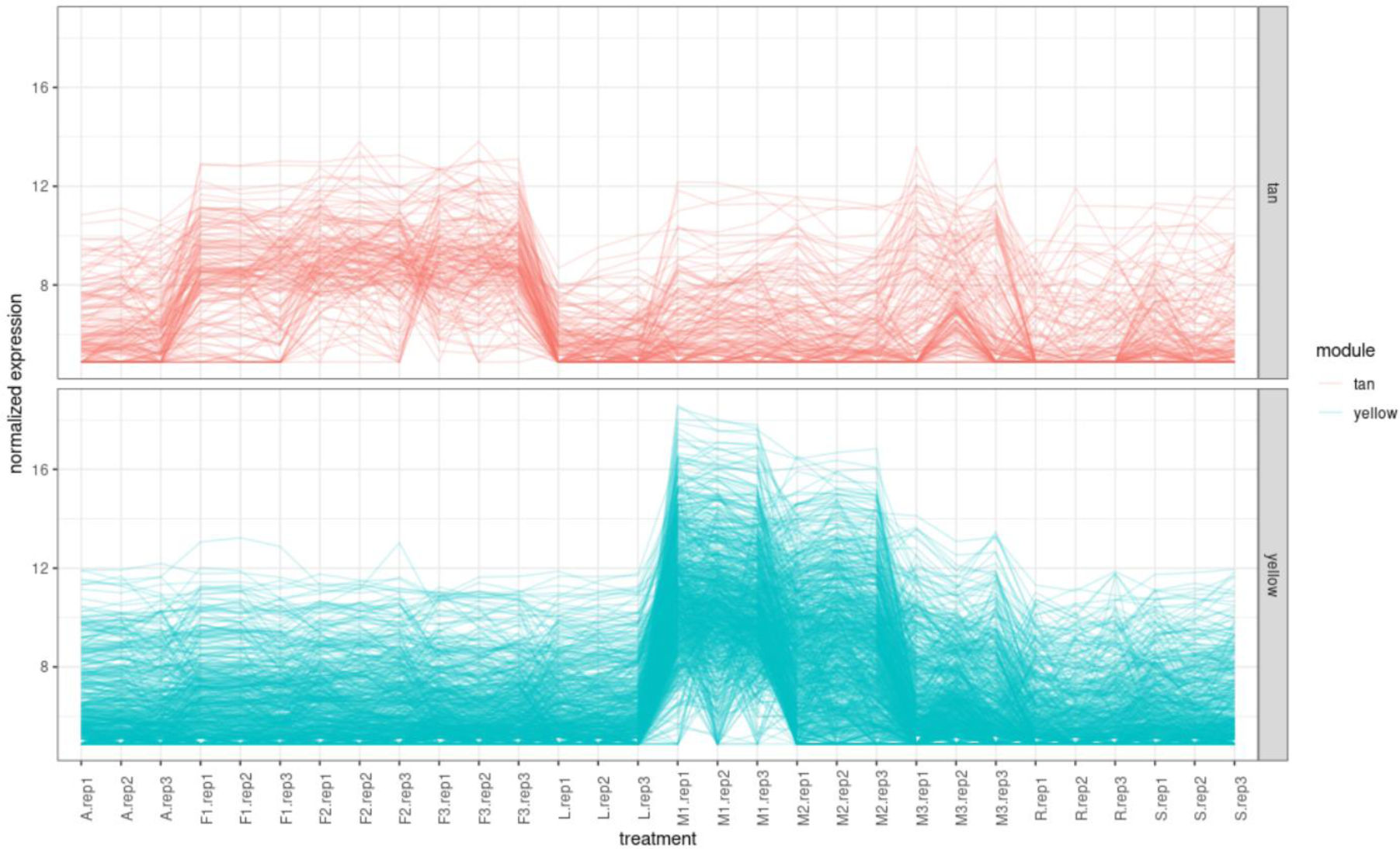
Modules of the interest. ‘tan’ module is positive for female cones (F1∼F3) and ‘yellow’ module is positive for male cones (M1∼M3).

We visualized the gene clusters in each of the selected modules (Fig. 5). ‘TRINITY_DN41963_c0_g1_i1’ encoding *AGAMOUS-like 6* (*AGL6*) (AT2G45650.1) and ‘TRINITY_DN2171_c0_g1_i4’ encoding *MYB DOMAIN PROTEIN 80* (*MYB80*) (AT5G56110.1) were identified as the hub genes of the tan and yellow module, respectively.

**Figure 5.**
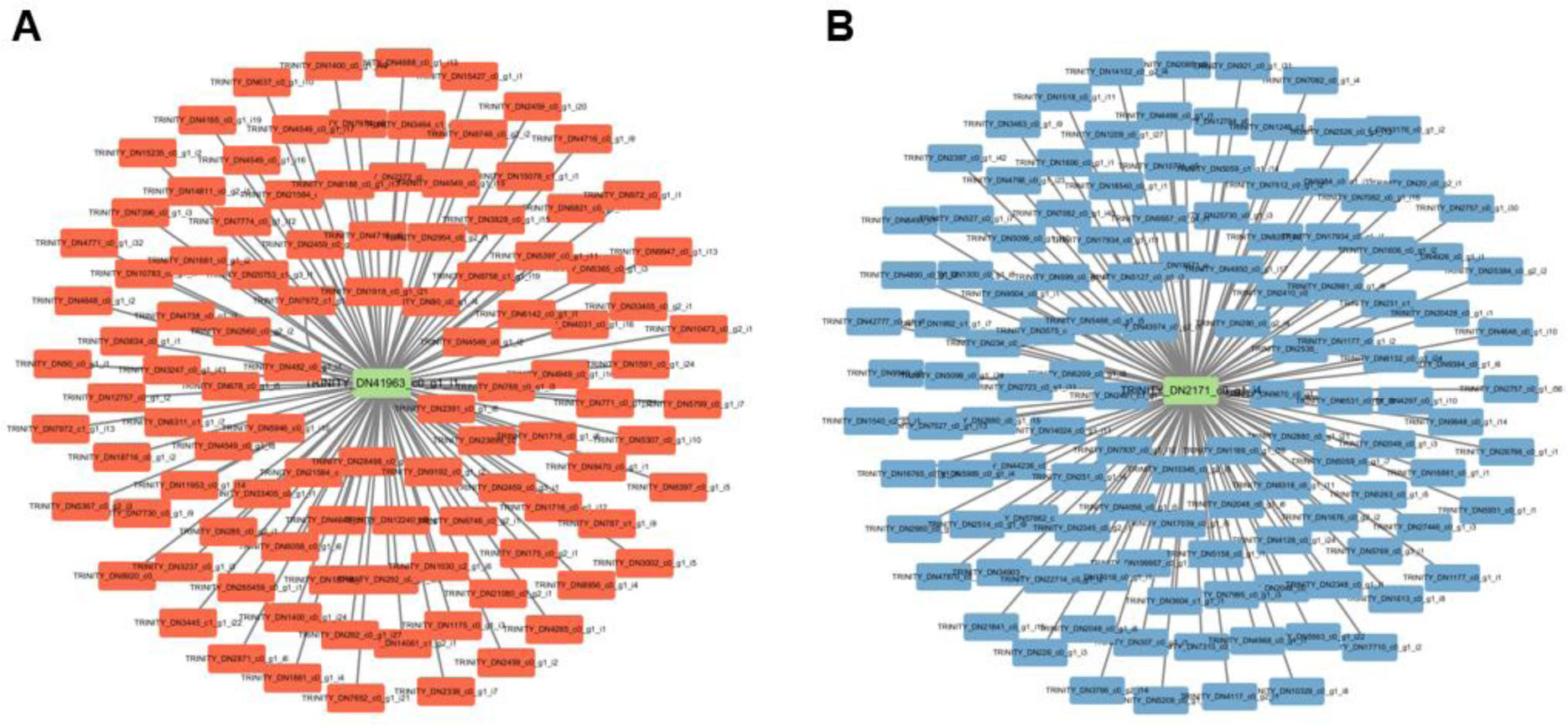
Co-expression gene network analysis. **A)** The tan module for female cones. **B)** The yellow module for male cones. Green rectangles represent the hub genes involved in reproductive development.

### Validation of RNA-seq data with qRT-PCR expression values

To confirm that the expression levels of transcriptomes were accurately measured using RNA-seq data in this study, total twelve DEGs were randomly selected and amplified by qRT-PCR. With a correlation coefficient of 0.74, we could decide that the relative expression profiles of the DEGs using RNA-seq platform was consistent with qRT-PCR approach (Fig. 6). In other words, it was judged that the results of this study, which were analyzed and interpreted by RNA-seq data, were reliable.

**Figure 6.**
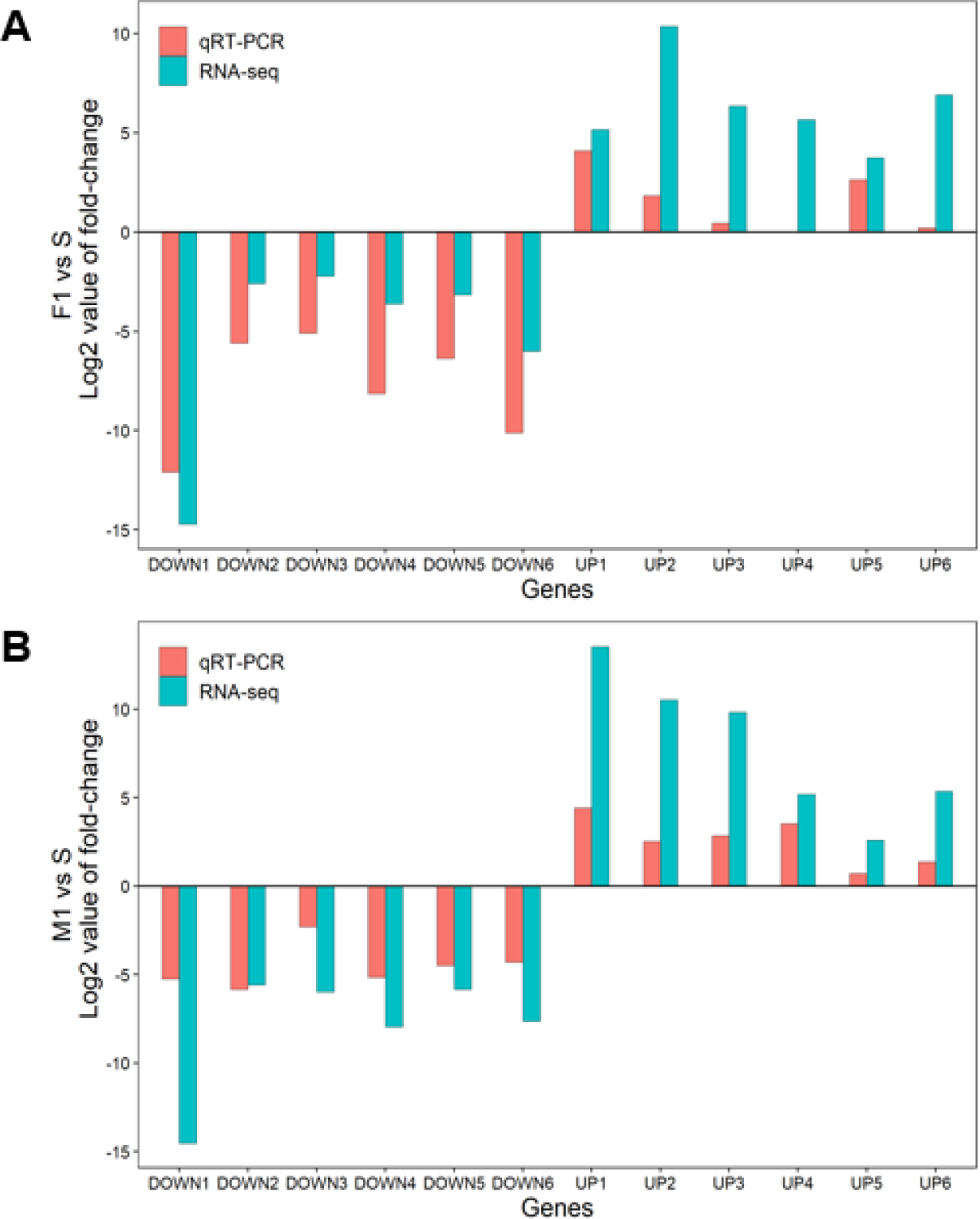
Validation of RNA-sequencing results by quantitative real-time PCR (qRT-PCR) using up- and down-regulated differentially expressed genes. **A)** Log_2_ values of fold-change between the early female cone (F1) and stem (S). **B)** Log_2_ values of fold-change between the early male cone (M1) and stem (S). Expression values are means for three biological replicates. DOWN, the down-regulated genes; UP, the up-regulated genes.

## Discussion

### Lignin biosynthesis and cell wall loosening are necessary for female cone development

Conifer species have sex-specific reproductive structures that are pollen-bearing male cones and ovule-bearing female cones (Rudall et al., 2011). Each developmental stage of the female and male cones of *Pinus densiflora* showed different gene expression characteristics. In the early stage of female cone development, lignin biosynthesis was active, and in the late stage, cell wall loosening acted predominantly. In *Pinus resinosa*, the thickening and lignification of cells are known to be necessary for the opening of cone scales (Dickmann and Kozlowski, 1969). In addition, cell walls of seed cones in the genus *Pinus* (*Pinus banksiana*, *P. nigra*, *P. ponderosa*, and *P. taeda*) have been reported to have unique chemical properties to provide resistance to decay (Thomas and Raymond, 1996).

Meanwhile, expansion or major cell wall disruption requires cell wall loosening, and it is reported that cell wall loosening of the mature pollen grains occurs at the surface of the stigma during pollen-stigma interaction (Carol et al., 1998). Therefore, it is presumed that lignin synthesis actively occurs in the early developmental stages of the female cones in *P. densiflora*, so that scale is opened, and in the later stage, pollen differentiation occurs by cell wall disruption after pollination and fertilization are performed. Through these processes, the female cones eventually become seed cones.

### Formation of pollen wall and syncytium is the major biological process in the early male cones

Metabolism of lipid and pectin, synthesis of lignin, and the formation of syncytium were predominant in the early stage of male cone development compared to the later stage. Syncytium refers to a multinucleated cell structure that appears because of the fusion of several single nucleated cells (Daubenmire, 1936; Lin et al., 2021). A major structural process involved in early pollen development is the syncytium formation of microspore mother cells (MMCs) or pollen mother cells (PMCs), followed by separation of each MMC to create microspores surrounded by callose walls (Shivanna et al., 1997). In addition, there is a specialized layer within anther, called tapetum, which is important for the development and nutrition of pollen grains, and then the tapetum also forms syncytium (Polowick and Sawhney, 1993; Clément and Audran, 1995).

In plants, the walls of mature pollen grains are composed of pectocellulosic intines containing pectin and cellulose components, and exines composed of sporopollenin, a lipid- and phenolic-based polymer (Scott et al., 2004; Morant et al., 2007). Lipid metabolism is essential for exine development, the outer pollen wall, and it has been reported that restricting lipid metabolism induces genic male sterility (GMS) during anther and pollen development in several species including *Arabidopsis*, rice, and maize (Shi et al., 2015; Wan et al., 2020). Through this, it was confirmed that in *P. densiflora*, MMC forms syncytium in the early stage of male cone development, and then pollen walls are created by metabolism of lipid and pectin along with lignin, a component of the plant cell wall.

### Different levels of phytohormone can influence the development of sex-specific cones in *P. densiflora*

Phytohormone plays an important role in the reproductive development of plants and is known to have a great influence on the cone growth in various gymnosperms. *Dacrydium pectinatum* maintains low level of gibberellin (GA), indole acetic acid (IAA), and abscisic acid (ABA) to promote seed germination in female cones, and the levels of GA and IAA fluctuated during the male cone development (Lu et al., 2021; Wang et al., 2022). In addition, GA treatment induced both female and male cones in Douglas-fir during long shoot development (Kong et al., 2008).

In this study, the sex-specific gene expression of the female and male cones of *P. densiflora* were shown. In particular, GA and IAA-related genes were highly expressed in the early female cones compared to the early male cones. In the case of female cones of *P. tabulaeformis*, the IAA content is highest in the early stage of development and gradually decreases thereafter, and it is reported that IAA defects induce female gametophyte abortion (Ren-yan and Cai-xia, 2005). Through this, it is thought that GA and IAA hormones are essential for the early reproductive growth of female cones in *P. densiflora*, and further studies on various hormones involved in the early development of both female and male cones are needed in the future.

We found hub genes that play an important role in the development of each reproductive organ by searching for gene clusters that are highly expressed in each of the female and male cones of *P. densiflora*. *AGAMOUS-like 6* (*AGL6*) and *MYB DOMAIN PROTEIN 80* (*MYB80*) were the hub genes of the female and male cones, respectively. *AGAMOUS* (*AG*) is a MADS-box family gene required for normal flower development in *Arabidopsis* and it confers the identity of floral organs such as stamens, carpels, and ovules and determines the floral meristem (Bowman et al., 1989; Yanofsky et al., 1990; Sieburth and Meyerowitz, 1997; Dreni and Kater, 2013).

*AGAMOUS-like 6* and *MYB DOMAIN PROTEIN 80* are the hub genes of female and male cone development of *P. densiflora*.

In ornamental plants, core regulatory networks that regulate the flower development of *Chrysanthemum morifolium* are explored and hub genes are discovered, providing information on the mechanism that determines flower shape (Ding et al., 2020). In addition, comparative transcriptive analysis in *Cymbidium nanulum* has found differentially expressed genes and candidate genes involved in flower development (Fu et al., 2022). In trees, there is also a study that performs transcriptomic analysis to find candidate genes for the double flower phenotype of *Prunus mume* (Zhu et al., 2022). However, different genetic factors are shown to affect reproductive growth in *Pinus* species.

*AGL6* is known to be expressed in the endothelial layer of the ovule adjacent to the growing female gametophyte (Schauer et al., 2009). In gymnosperms, the expression of *AG* orthologous genes has been studied in several species. *TcAG* of *Taxus chinensis* var. *mairei* was expressed in both developing female and male cones (Liu et al., 2018), and *AG* orthologous genes were also expressed during the development and ripening of the fleshy fruit-like structures in *Ginkgo biloba* and *Taxus baccata* (Lovisetto et al., 2012). Therefore, it can be suggested that *AGL6* plays a central role regulating a transcriptional network during the development of the female cones in *P. densiflora*.

In *A. thaliana*, the regulation of tapetal programmed cell death (PCD) and pollen formation depend on a transcription factor MYB80, formerly known as MYB103. Tapetum is the inner cell layer that surrounds growing microspores within the anther and is crucial for providing nutrients and structural elements during the microspore development. Pollen maturation causes PCD in the tapetum, providing several chemicals necessary for the formation of pollen, including release of microspores and development of pollen walls (Parish and Li, 2010; Phan et al., 2011; Vardar and Ünal, 2012; Verma, 2019).

There have been few direct studies related to the expression of *MYB80* in male cones of gymnosperms, but it has been reported that some downstream genes of *MYB80* are important for male cone development. Several downstream genes including a pectin methylesterase, a glyoxal oxidase, and an A1 aspartic protease, which are all direct targets of *MYB80* in *Cryptomeria japonica*, are reported to be necessary for the male cone fertility (Wei et al., 2021). In view of these facts, *MYB80* is assumed to play a major role regulating a transcriptional network during the development of the male cones in *P. densiflora*.

In conclusion, identifying main biological factors and key genes involved in reproduction is important for reducing the long breeding cycle of forest trees. Here, we investigated major biological processes and hub genes involved in the cone development of *P. densiflora* based on comparative transcriptomic analysis. These results can facilitate further efforts to regulate the onset of reproduction phase and modify cone production in seed orchards of *Pinus* species.

## Materials and Methods

### Plant materials and sample collection

Tissues were sampled from three *P*. *densiflora* Sieb. & Zucc. adult trees grown in Chilbosan experiment forest of Seoul National University (Suwon-si, Gyeonggi-do, Korea) between March and May 2022. Three stages of reproductive tissues during the cone developmental period were sampled (Supplemental Fig. S5): early female cone (F1) and early male cone (M1); mid female cone (F2) and mid male cone (M2); late female cone (F3) and late male cone (M3). Female cones began to be collected a week later than male cones.

In addition, vegetative tissues were collected once from the same trees and the following tissues were sampled: from the tip of the shoot, apex (A), young needles (L), and young stem (S); fine root (R). Each tissue type was represented by three biological replicates and the sample collection is detailed in Supplemental Table S2. All plant material was flash-frozen in liquid nitrogen and subsequently stored at −80° until RNA extraction.

### RNA extraction, library preparation, and Illumina sequencing

Total RNA of plant tissues was extracted using IQeasy™ plus Plant RNA Extraction Mini Kit (iNtRON, Seongnam-si, Gyeonggi-do, Korea). The RNA quantity and purity were assessed using Agilent 4200 TapeStation (Agilent Technologies, Santa Clara, CA, USA). For library preparation, 500 ng of total RNA from each sample with RNA integrity number (RIN) ≥ 8.0 was used. 30 cDNA libraries (> 20 nM) were sequenced on the Illumina HiSeq2000 platform (Illumina, San Diego, CA, USA), producing paired-end reads of 100 bp. Library preparation and sequencing were conducted at the LabGenomics (Seongnam-si, Gyeonggi-do, Korea). The sequencing results of each library can be found in Supplemental Table S3.

We checked the quality of raw reads using FastQC (v0.11.9) (https://www.bioinformatics.babraham.ac.uk/projects/fastqc/). Adapter sequences were trimmed from the reads, and the read sequences were filtered for quality and minimum length using Trimmomatic (v0.36) (Bolger et al., 2014).

### *De novo* transcriptome assembly and differential expression analysis

After trimming and filtering, all reads were assembled for generating a single *de novo* reference transcriptome using Trinity software (v2.11.0) (Grabherr et al., 2011). Sequences satisfying an open reading frame (ORF) of 100 amino acids or more were then extracted using TransDecoder (v5.5.0) (https://github.com/TransDecoder/TransDecoder). To remove isoforms, we subsequently clustered the entire sequences using CD-HIT (v4.8.1) (Li and Godzik, 2006). The longest sequence among 99% similar sequences in each cluster was extracted and these were used as the representative transcript set for downstream analysis.

To quantify gene expression levels, clean reads were mapped on the assembled transcriptome of *P. densiflora* by RNA-Seq by Expectation-Maximization (RSEM) method (Li and Dewey, 2011). The expression level of each transcript was normalized by Trimmed Mean of *M*-values (TMM). Differentially Expressed Genes (DEGs) meeting criteria of at least four-Fold Change (FC) and a *p*-value less than 0.001 were identified using edgeR package (v3.34.1). Especially, we extracted DEGs from four pairwise comparisons: female cone early vs female cone late, female cone early vs male cone early, and male cone early vs male cone mid. These DEGs were analyzed for the functional annotation and GO enrichment analysis as described in the Gene annotation and GO enrichment analysis section.

### Gene annotation and GO enrichment analysis

BLASTp (https://blast.ncbi.nlm.nih.gov/Blast.cgi) searches of the TAIR10 protein databases (https://www.arabidopsis.org/index.jsp) were conducted for gene annotation of DEGs in each comparison. Gene Ontology (GO) enrichment analyses were performed using DAVID (https://david.ncifcrf.gov/home.jsp) with an *Arabidopsis thaliana* background. GO terms with an adjusted *p*-value (Benjamini and Hochberg, 1995) below 0.05 were considered to be significantly enriched.

### Data processing for co-expression gene network analysis

Weighted Gene Co-expression Network Analysis (WGCNA) was performed to identify the target modules specific to female and male cones of *P. densiflora*. We used WGCNA (v1.72-1) R package using the total of count data matrix as an input file to obtain a list of clustered genes and weighted gene correlation network files as the output files (Zhang and Horvath, 2005; Langfelder and Horvath, 2008). First, the counts were normalized with DESeq (v1.32.0) and 95 quantile expression data was used for downstream analysis. Then, the weighted adjacency matrix is defined and the soft threshold power 10 was selected because the scale free topology fitting index and mean connectivity may appear to reach a plateau (Supplemental Fig. S6).

Each module for female cones and male cones was selected and the correlation networks of the modules were visualized using Cytoscape (v3.10.0). We only used genes which are involved in reproductive development and with a correlation *p*-value below 0.05 among the clustered genes in each module for the hub gene identification.

### Quantitative real-time PCR

Quantitative real-time PCR (qRT-PCR) was used to verify basic expression levels of a subset of genes in the assembled transcriptomes of *P. densiflora*. Three types of tissues (F1, M1, S) with three biological replicates were used for qRT-PCR. Twelve DEGs were randomly selected: six up-regulated and six down-regulated in both F1 and M1 compared to S. We used 18s rRNA sequence in the assembled transcriptomes of *P. densiflora* as a reference gene to normalize the expression of the investigated genes.

Quantity of 100 ng of total RNA per sample was used to synthesize cDNA using a ReverTra Ace® qPCR RT Master Mix (TOYOBO, Kita-ku, Osaka, Japan). The synthesized cDNA was diluted with nuclease-free water, and 100 pg of cDNA per sample was used for final concentration. The amplification process was performed in a CFX Connect™ Real-Time System thermal cycler (Bio-Rad, Hercules, CA, USA) using ExcelTaq™ 2X Fast Q-PCR Master Mix (SYBR, no ROX) (SMOBIO, Roswell, Georgia, USA) under the following conditions: 95° for 30 seconds, then 40 cycles of 95° for 15s and 60° for 30s, followed by a melt curve stage increasing from 65° to 95° at a rate of +0.1° s^-1^. The relative expression levels were calculated with Bio-Rad CFX Maestro software (v2.2) using the 2^-ΔΔCt^ method. All primers were made using Primer3 (v4.1.0) (https://bioinfo.ut.ee/primer3/), and the information of the primers is detailed in Supplemental Table S4.

## Acknowledgments

We would like to thank Professor Sangrea Shim from Kangwon National University for his valuable comments and suggestions in the data analyses.

## Author contributions

D.L. and Y.K. devised and designed the research plans. Y.K. and K.K. supervised the experiments. D.L. and Y.K. performed experiments and analyzed the data. D.L. wrote the manuscript with the help of K.K..

## Supplemental data

The following materials are available in the online version of this article.

**Supplemental Figure S1.** Sample correlation results.

**Supplemental Figure S2.** Number of Differentially Expressed Genes between tissue samples of *Pinus densiflora*.

**Supplemental Figure S3.** Hierarchical clustering dendrogram of gene modules with different colors identified by Weighted Gene Co-expression Network Analysis.

**Supplemental Figure S4.** Relationship of tissue types and module eigengenes.

**Supplemental Figure S5.** Structure of reproductive tissues of *Pinus densiflora*.

**Supplemental Figure S6.** Different soft-thresholding powers.

**Supplemental Table S1.** Gene Ontology enrichment analysis results of selected modules for the female and male cones in *Pinus densiflora*.

**Supplemental Table S2.** Sampling information of the reproductive and vegetative tissues of *Pinus densiflora*.

**Supplemental Table S3.** Statistics of raw sequencing data.

**Supplemental Table S4.** Oligonucleotide primers used for validation by quantitative real-time PCR.

## Data availability

All data are available within the manuscript and its Supplemental data.

